# Mixing selects for predation resistance in lab-evolved communities of bacterial prey and social predator *Myxococcus xanthus*

**DOI:** 10.1101/2023.03.21.533597

**Authors:** Saheli Saha, Bhoomika Bhat, Jessica Mellicent Laloo, Akshit Goyal, Samay Pande

## Abstract

Population mixing, and transmission modes strongly influence social interactions. However, the impact of repeated mixing on the evolutionary dynamics of microbial predator-prey interactions remains underexplored^1,2^. Hence, we conducted a laboratory evolution experiment with bacterial predator-prey communities under two transfer regimens: repeated mixing (horizontal transfer) versus no mixing (vertical transfer). For this, *Myxococcus xanthus* served as the generalist predator^3,4^ and *Escherichia* coli as prey. We show that prey populations from vertical regimen were less resistant to predation than the ones from horizontal regimen. This was because prey isolates in the vertical treatment showed varying resistance levels relative to ancestors, while those in the horizontal treatment demonstrated less variation. Moreover, analysis of prey populations over evolutionary time revealed that the populations from horizontal treatment showed increasing levels of resistance to predation over time, whereas the ones from vertical treatment did not show similar trend. The differences in the outcomes of the two treatments was because the variants better at intraspecies competition, can only be maintained in the vertical treatment, whereas in horizontal treatment the benefits of superior intraspecies competitive fitness are nullified because of population mixing, as predicted by mathematical modeling approach. These predictions were empirically confirmed. Moreover, contrary to our expectations, predatory efficiency of evolved *M. xanthus* isolates was either the same or lesser than the ancestor. Together, we demonstrate that mixing affects the evolution of prey bacteria but has little effect on the hunting ability of the social predator *M. xanthus*.

## Introduction

The influence of mixing on the evolution of microbial interactions and their effects on community assembly has garnered significant interest in recent times ^1,2^. This interest seems justified, given that the mixing has been shown to strongly affect the evolution of antagonistic as well as synergistic interactions within populations (i.e., groups of individuals of single species) ^1,2,5–9^. Microbial communities in nature are frequently mixed because of human activities, abiotic, and biotic factors ^10–12^. However, despite the prevalence of mixing events, we have limited understanding of the effects of repeated mixing on the evolution of microbial interactions both in the context of simple communities such as predator-prey interaction, and complex multispecies communities ^1,2,13^.

Evolutionary trajectories of antagonistically interacting species, like host-parasite and predator-prey systems follow the Red Queen (RQ) dynamics in which interacting partners coevolve in an evolutionary arms-race ^14–16^. The RQ hypothesis has gained significant prominence over the years and has been proposed to explain the evolution of sex, antagonist-driven emergence of diversity, symbiosis, and antimicrobial resistance ^8,17,18^. Originally proposed by Van Valen (1973) ^19^, the Red Queen hypothesis is intentionally focused only on the influence of the biotic factor. However, in nature, a combination of both biotic and abiotic factors will likely influence the outcome of antagonistic evolution. Therefore, extensions of this theory that incorporate both the effects of biotic and abiotic factors, such as diversity, migration, and changes in environmental conditions have attracted considerable attention^7^. For example, studies using host-parasite interactions suggest a significant effect of population mixing on the Red Queen dynamics between co-evolving antagonists ^18,20^. While predator-prey coevolutionary dynamics has been the focus of multiple studies in the past ^21–24^, surprisingly the influence of repeated mixing, across habitats or even ecosystem, on the predator-prey coevolutionary dynamics is relatively less explored among microbes ^1^.

Here we asked whether repeated mixing of distinct microbial predator-prey consortia will affect their coevolutionary dynamics. For within-species interactions, theory and empirical evidence suggest that the coevolutionary dynamics without mixing events are likely to promote synergistic interactions^25^. In contrast, repeated mixing of microbial consortia, that are growing independent of each other are likely to promote antagonism ^25^. Hence, we predicted that for interspecies interactions (such as predator-prey) too, repeated mixing will result in the evolution of predators that show better predatory performance relative to the ones that evolved in communities that did not experience repeated mixing events. Similarly, prey populations that experience repeated mixing events will evolve to be more resistant to predation than when predator-prey communities are propagated independent of mixing events. This is because of differences in the partner-fidelity between the two treatments. We also hypothesized that the populations from mixed (horizontal) and unmixed (vertical) treatments would follow distinct evolutionary strategies. This is because the mixing events will increase the supply of mutation which might affect the fixation probabilities (by selecting larger effect mutations) because of changes in the clonal interference. To test these hypotheses, bacterial predator-prey communities of *M. xanthus* and *E. coli* were propagated in a lab evolution experiment. *M. xanthus* is a generalist bacterial predator that kills prey bacteria by both contact-dependent and contact-independent mechanisms ^4,26–30^. This includes the production of enzymes ^27,31^, antibiotics ^3^, and secretion systems ^28,30^. In the presence of *M. xanthus,* coevolving *E. coli* populations have been shown to diversify and maintain predation-resistant mucoid variants along with the predation-sensitive types over the evolutionary timescale ^24,32^. Further, as expected in a winnerless Red Queen dynamic, *M. xanthus* evolves higher predatory performance^24^.

Here, we demonstrate that coevolution with a social microbial-predator results in an asymmetric arms-race where only the prey continues to evolve better survival strategies over evolutionary time in response to predation. Further, we show that the mixing of predator-prey populations during each transfer results in the evolution of resistance to predation. Whereas prey populations that do not experience mixing events maintained greater diversity in prey phenotype based on their varying degree of resistance to predation, indicating maintenance of diverse strategies. Most interestingly, evolved populations of predators from both regimens show the emergence of within-population diversity which harbors genotypes with similar predatory performance as their ancestors and the genotypes with reduced predatory performance relative to their ancestors. The mathematical modelling approach revealed that the differences in the outcomes between the two treatments are because of changes in the clonal interference caused by mixing events. These predictions were further confirmed by performing a laboratory evolution experiment.

## Results

### Repeated mixing results in increased resistance to predation in the evolved prey population

The evolution experiment designed to test the effect of mixing on the evolution of prey and predator populations involved repeated mixing of predator-prey consortia every fourth day (Supplementary figure S1) for eighty days. Whereas control populations were propagated without mixing. To perform these experiments glucose was used as a carbon source. Since *E. coli* can use glucose as a carbon source but *M. xanthus* cannot, predation on *E. coli* population was the only source of nutrients for *M. xanthus*. The evolved communities after sixty-four days of serial transfers were analysed for the changes in the resistance to predation in the evolved *E. coli* and changes in the predatory performance of evolved *M. xanthus*.

To test the effect of repeated mixing, we measured the growth of evolved (after sixteen rounds of propagation in the lab evolution experiment) and ancestral *E. coli* populations when cocultured with ancestral *M. xanthus* in conditions identical to the ones during the evolution experiment. This measure accounts for the growth and survival of prey in the presence of *M. xanthus*. In line with our expectations, *E. coli* populations from the horizontal regimen on average exhibit better growth in the presence of predators than their ancestors (1.09 log-fold relative to ancestor, one sample t-test, *p^[FDR^ ^corrected]^* < 0.001). Whereas growth of populations from vertical treatment did not show a significant difference relative to the ancestor (1.03 log-fold relative to ancestor, one sample t-test, *p^[FDR^ ^corrected]^* > 0.05). Thus, *E. coli* population from vertical treatment were in general less resistant to predation than the ones from horizontal treatment (Figure 1A, independent sample t-test, *p* = 0.057). However, one of the four vertically coevolved populations (population 3) did evolve higher levels of resistance to predation, which was comparable to horizontally transmitted lines (Figure 1A, Supplementary figure S2). Together, vertically propagated prey populations seem to evolve different mechanisms of survival in the presence of predators, which in most cases increases the sensitivity of the prey populations to predators.

**Figure 1:**
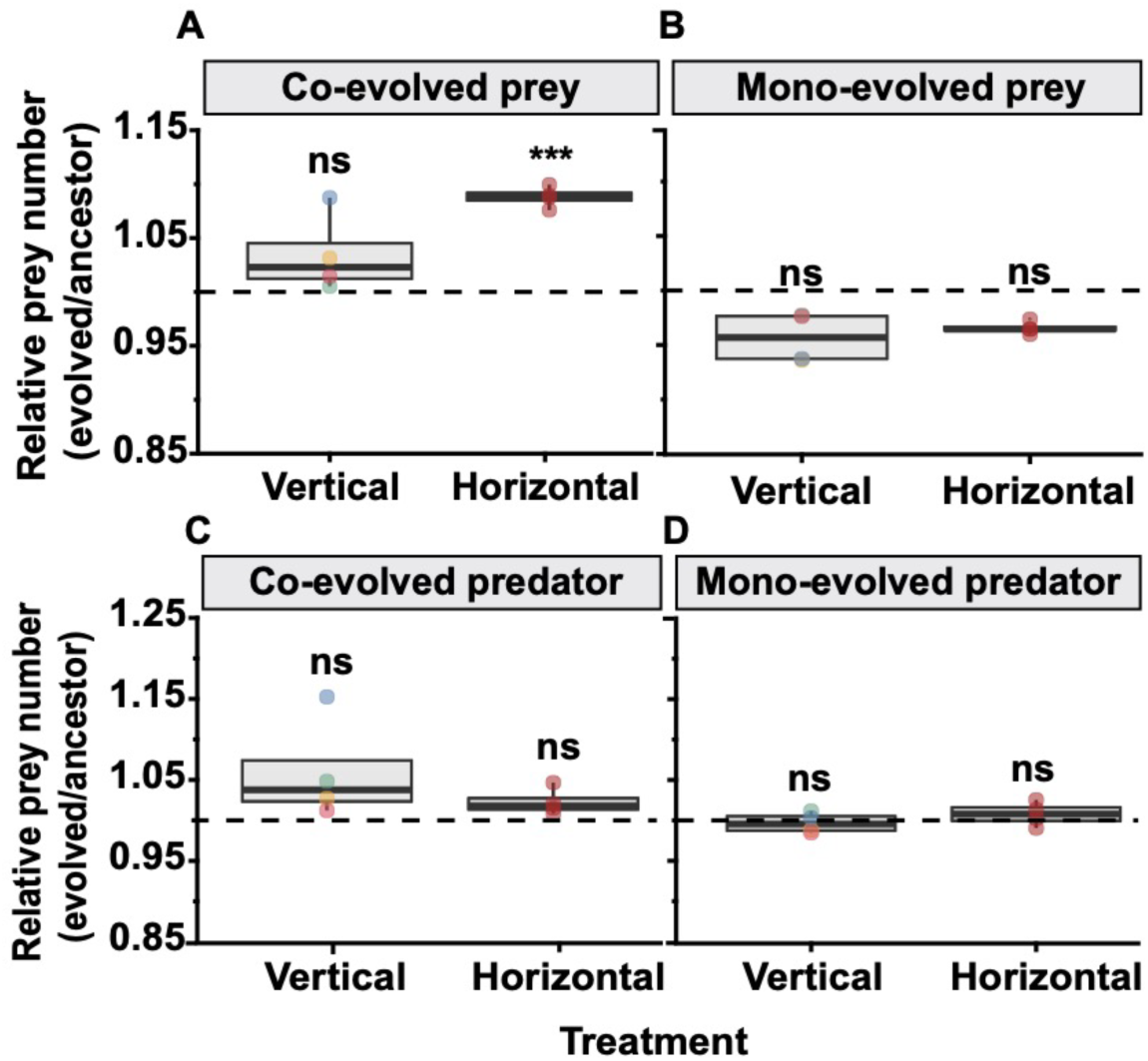
Effect of selection regimen on the evolution of resistance to predation in prey and predatory performance of *M. xanthus* populations. (A and B) Growth of evolved *E. coli* relative to the growth of ancestral *E. coli* in the presence of ancestral predator is shown. (A) Prey populations coevolved in the vertical transfer regimen were more susceptible to predation than the populations that evolved in the horizontal transfer regimen (paired t-test for differences between vertically evolved *E. coli* relative to horizontally co-evolved populations, p*** < 0.001). (B) Whereas mono-evolved populations showed no significant differences both between treatments as well as relative to the ancestors (Wilcoxon-test for differences between vertically and horizontally mono-evolved populations, *p* > 0.05, n = 4). Color of dots in the vertical regimen represents distinct lineages. (C) Predator populations coevolved in vertical and horizontal treatment exhibited similar predatory performance, which was not significantly different from their respective ancestors. Relative numbers (evolved/ancestor) are log(CFU/ml) of ancestral *E. coli* when cocultured with evolved predator population relative to their respective ancestor (one sample T-test for differences between coevolved predator population and their ancestors, *p* > 0.05, n=4). (D) Mono-evolved predator populations showed no significant differences both between treatment as well as from ancestors *p* > 0.05, n=4). Relative numbers (evolved/ancestor) in log (CFU/ml) of ancestral *E. coli* when cocultured with evolved predator population relative to their respective ancestor. Different color of dots in the vertical regimen represents distinct lineages.

Differences between the vertically coevolved and horizontally coevolved prey populations can emerge in response to the presence of predators or as an outcome of adaptation to abiotic factors. However, since mono-evolved *E. coli* populations (propagated in the absence of *M. xanthus,* Supplementary figure S1C and S1D) from the vertical as well as horizontal regimen showed no difference in the overall levels of resistance to predation (Figure 1B and Supplementary figure S2, one sample Wilcoxon test, *p* > 0.05), we concluded that the differences between vertical and horizontal *E. coli* populations propagated along with *M. xanthus* were not the result of adaptation to abiotic factors. Together, these results demonstrate a strong effect of the transfer regimen on the evolution of the prey population.

### Evolution of reduced predatory performance in co-evolved *M. xanthus* populations

It is difficult to reliably measure the growth of *M. xanthus* in our experimental setup (See methods). Hence, to assess changes in the evolved predator populations, the growth of ancestral prey was used as a measure of the predatory performance of *M. xanthus*. This measure of predation is a combination of growth and death of *E. coli* in the presence of the predator. Contrary to our expectations and previously reported findings ^24^, we observed that the ancestral *E. coli* was able to grow to a similar density in the presence of evolved as well as ancestral *M. xanthus* populations (Figure 1C, and Supplementary figure S3, one sample t-test *p* > 0.05). Further, no significant differences between coevolved *M. xanthus* populations (after sixteen transfers) from horizontal and vertical regimens (Figure 1C, independent sample t-test for differences between prey growth in the presence of evolved predators, *p* > 0.3) were observed. Together, the results suggest that the predator populations were probably under relatively weaker selection to evolve enhanced hunting ability. Thus, adaptive response other than the increased predatory performance as measured in our experiments might have been selected in our experiments.

It is expected that the evolved populations are internally diverse and hence population-level data might not be reflective of the intra-population diversity. Hence, we measured the predatory performance of distinct clones isolated from evolved *M. xanthus* populations. To do so, four *M. xanthus* clones were randomly isolated from each of the replicate lines from the vertical regimen and sixteen clones were isolated from the horizontal regimen. Surprisingly, a minority of the clones from both the transfer regimens showed reduced hunting performance relative to the ancestors (Supplementary figure S3, ANOVA and SNK *post hoc* test). Whereas the majority showed no difference.

However, genomic analysis of the evolved clones revealed mutations specific to the co-evolved clones that were absent in the ancestors. Most of these mutations were either located in the genes that putatively code for non-ribosomal peptide synthetases (NRPS) or transporter proteins (Supplementary table T4). Previous reports have demonstrated that NRPS is involved in the synthesis of antimicrobial peptides and hence is a major component of predatory behavior of *M. xanthus* ^33,34^. Thus, mutation in this locus is in line with the selection pressure during the evolution experiment. While in the bad predators (clones with lower predatory efficiency) most consistent mutations were in the genes that putatively code for chemotaxis proteins (Supplementary table T4). Mutations in the chemotactic pathways like in the CheA gene, which is a part of the Che3 cluster of chemotaxis proteins in *M. xanthus*, also explain reduced swarming efficiencies of the *M. xanthus* clones that have evolved reduced predatory efficiency (Supplementary table T4). The evolution experiments were conducted in a spatially structured environment, thus *M. xanthus* cells might need the ability to find the next patch of prey populations. Thus, chemotactic mutants that cannot swarm towards the prey show lower prey-killing efficiencies. Interestingly, despite the similarities in the predatory efficiencies of the predators from both regimens, the overall mutations were more similar in the horizontally coevolved *M. xanthus* isolates, compared to the vertically coevolved ones (Supplementary table T4). These results therefore strongly suggest that the predators’ populations have also evolved differentially. However, the differential evolution has not affected the prey-killing efficiency of the predator population.

### Coevolved *E. coli* clones from vertical regimen exhibit large within-population differences in their sensitivity to *M. xanthus* predation than the ones from horizontal treatment

Since evolved prey populations from the two treatments were significantly different from each other, we further tested whether the population level differences were representative of the phenotypes of distinct clones within each population. To do so, four *E. coli* clones were isolated from each of the vertically propagated lines and sixteen clones were isolated from the horizontally evolved population. We observed significant variation among the isolates derived from vertically coevolved populations than between clones isolated from horizontal transfer regimen (Supplementary figure S4, average variance among vertical clones = 0.414, average variance across horizontal clones = 0.130, independent sample t-test for differences between variance *p* < 0.02). Importantly, out of sixteen clones from the vertical regimen, five clones seem to have higher resistance and four clones seem to show lower resistance to predation relative to ancestors (Figure 2 and Supplementary figure S4). Whereas all clones from the horizontal regimen were more resistant to predation than the ancestors. The evolution of *E. coli* strains sensitive to *M. xanthus* predation in vertically evolved populations suggested that the evolution of resistance to predation might not be the only adaptive strategy. This was indeed the case, as pairwise competition experiments between evolved *E. coli* clones and their respective Kanamycin-marked fitness-neutral ancestors (Supplementary figure S8) revealed that irrespective of their sensitivity to *M. xanthus* predation, most coevolved clones showed higher competitive fitness than ancestors in the presence of *M. xanthus* (Figure 3, one sample Wilcoxon test *p^[FDR^ ^corrected]^* < 0.0005). Moreover, the fitness of co-evolved clones was in general significantly higher in the presence of *M. xanthus* than in the absence of it (Figure 3, Kruskal Wallis and Dunn’s *post hoc* test).

**Figure 2:**
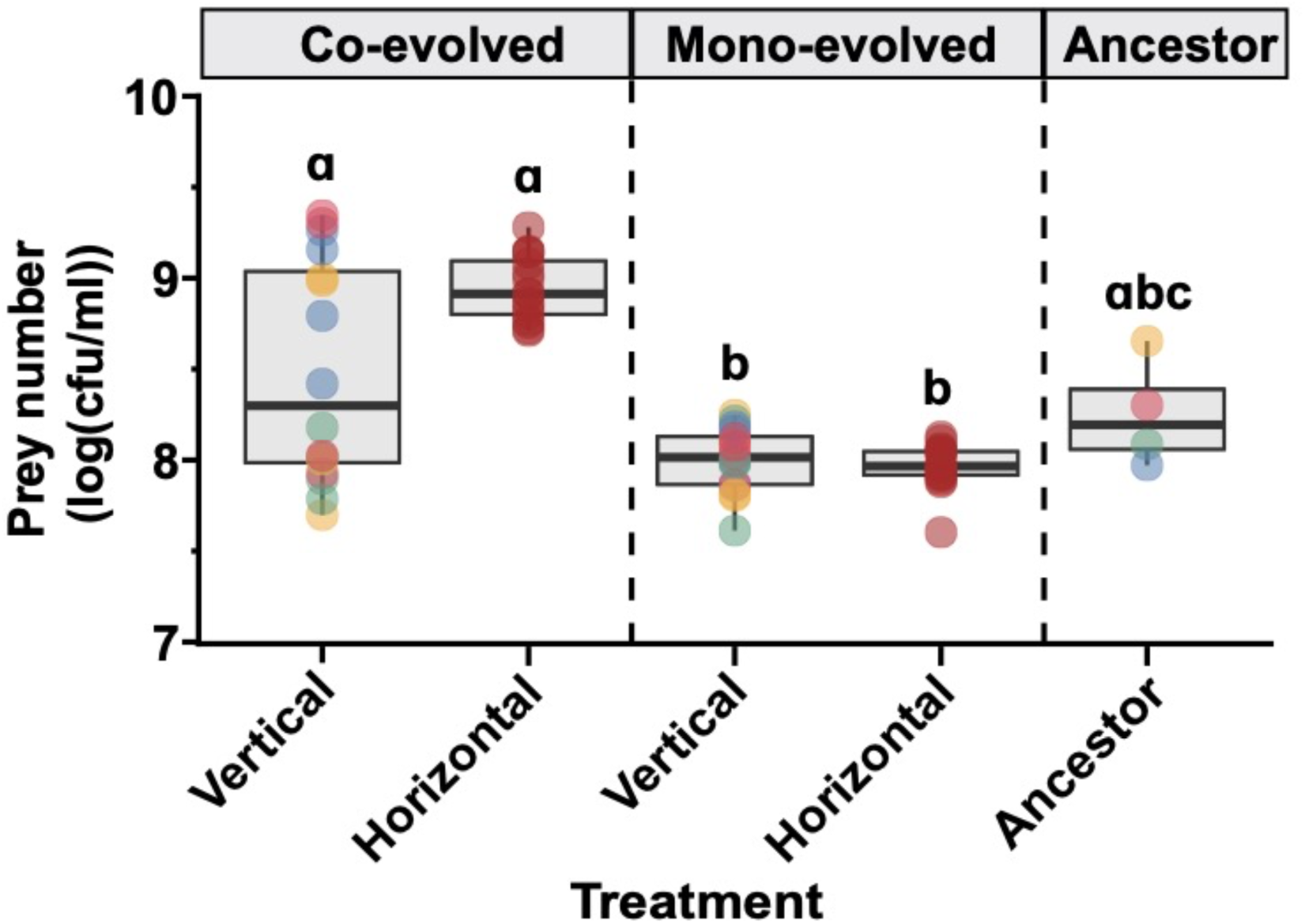
Effect of the selection regimens on the evolution of resistance to predation in prey clones. Survival of evolved and ancestor *E. coli* clones is shown as log(CFU/ml) of prey numbers when cocultured with the ancestral *M. xanthus.* Letters indicate significant difference between groups (Kruskal-Wallis, Dunn’s post hoc test, n =3 for each clone studied). Different color of dots in the vertical and ancestral regimen represents a distinct lineage that started from the ancestor shown in the same color.

**Figure 3:**
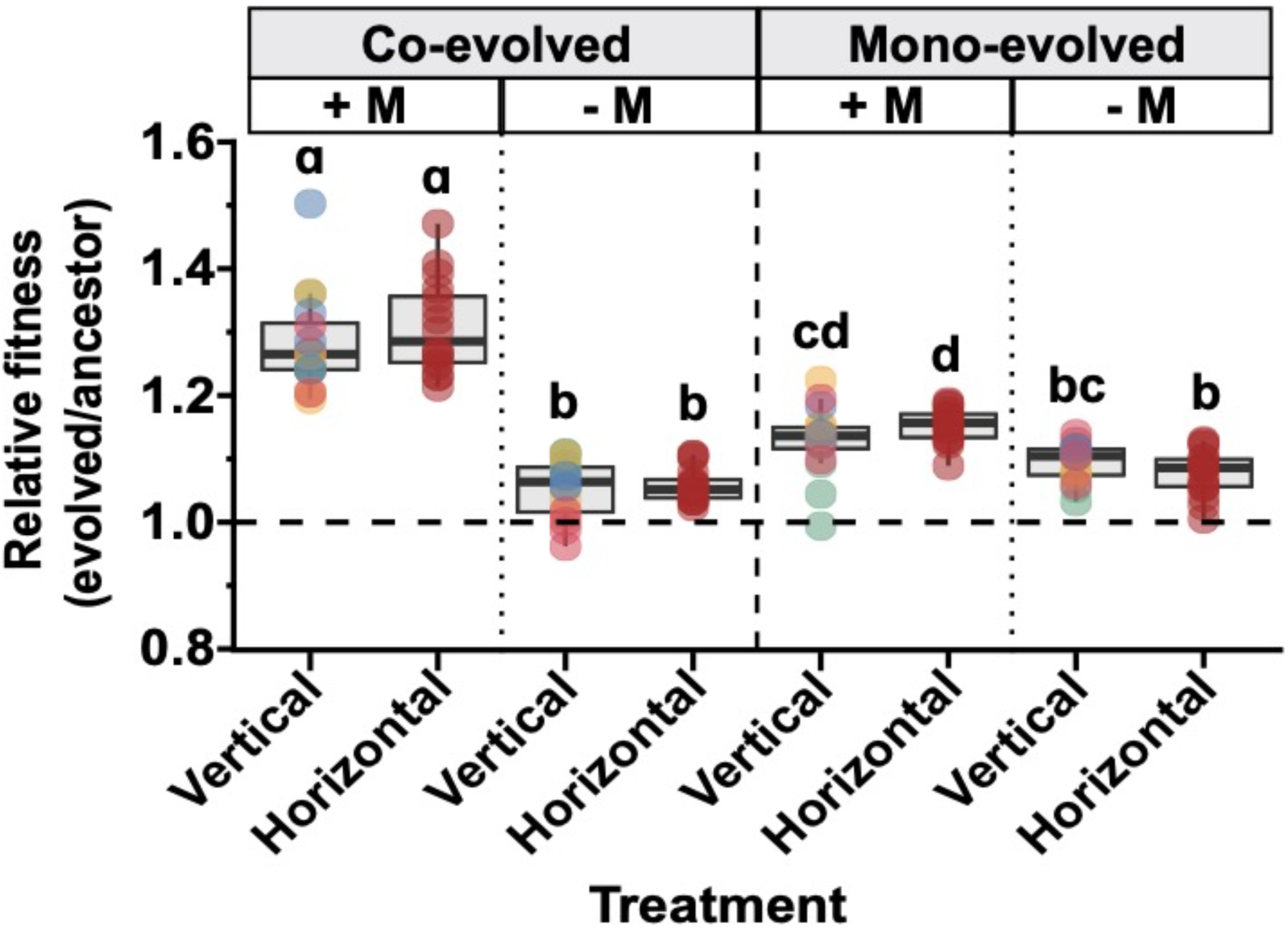
Effect of the selection regimens on the competitive fitness of prey clones relative to their ancestors. Relative fitness of evolved *E. coli* clones (dots) is shown. All groups are significantly different from 1.0 (one sample Wilcoxon test for differences between average relative-fitness of all clones relative to 1.0, *p^[FDR^ ^corrected]^*< 0.01, n = 3), and letters indicate significant differences between groups (Kruskal-Wallis, Dunn’s post hoc test for differences among four treatments). Different color of dots in the vertical treatment represents distinct lineages.

Similar experiments with strains from mono-evolved lines of *E. coli* showed that all the clones had lower resistance against *M. xanthus* than their ancestors (Figure 2, an average of prey survival of mono-evolved clones was 1.03-fold lower than ancestors, one sample Wilcoxon test, p < 0.0003). Further, we did not detect differences in the relative fitness of mono-evolved *E. coli* in the presence of *M. xanthus* than in its absence (Figure 3, Kruskal Wallis, Dunn’s *post hoc*).

Further analysis of the data from predation experiments (Figure 2) and competition experiments (Figure 3) revealed a positive correlation between resistance to predation and relative fitness for vertically coevolved clones of *E. coli* (Supplementary figure S5, Pearson’s correlation coefficient (r) = 0.6, p < 0.019), but not for *E. coli* coevolved in the horizontal regimen. Thus, though the evolution of high levels of resistance to predation is not essential for competitive advantage against ancestors, strains with higher resistance will generally have higher fitness than the ones with lower levels of resistance to predation.

Genomic analysis of the evolved and ancestral prey clones from both coevolved and mono-evolved treatments revealed several mutations that are specific to coevolved prey clones (Supplementary table T3). Amongst these, insertions at different locations in and after fliPQR operon in RcsB binding site were most common ^35^. This binding site is important for the regulation of RcsA gene that is involved in Rcs-phosphorelay^35^. Previous studies have demonstrated a diverse set of functions for Rcs signal transduction system that include regulation of motility, capsule formation and virulence ^35–37^. A detailed understanding of how these mutations affect the sensitivity of prey bacteria in our experiments will require exhaustive analysis that is not part of the current manuscript. Further, a higher genetic diversity was observed in the vertically coevolved prey, that is consistent with their phenotypic diversity. Whereas the horizontally coevolved prey was more genetically similar with comparable mutations in the majority of the clones.

### Mathematical modelling reveals prey evolution in horizontal treatment is mainly driven by adaptation to the presence of predator rather than intraspecific competition between prey

Together, our experiments demonstrate that horizontally propagated populations, which were mixed, only evolved predator avoidance, while vertically propagated ones, with no mixing, evolved both predator avoidance and prey competitive ability (Figure 2 and Figure 3). To explain why mixing suppressed prey competitive ability and amplified predator avoidance, we formulated a simple mathematical model of our communities. Briefly, our model partitioned each growth cycle into two phases: an initial period of prey growth, and a later period of predator growth and prey death (Figure 4A), consistent with experimental data of the dynamics within a growth cycle (Supplementary figure S6).

**Figure 4:**
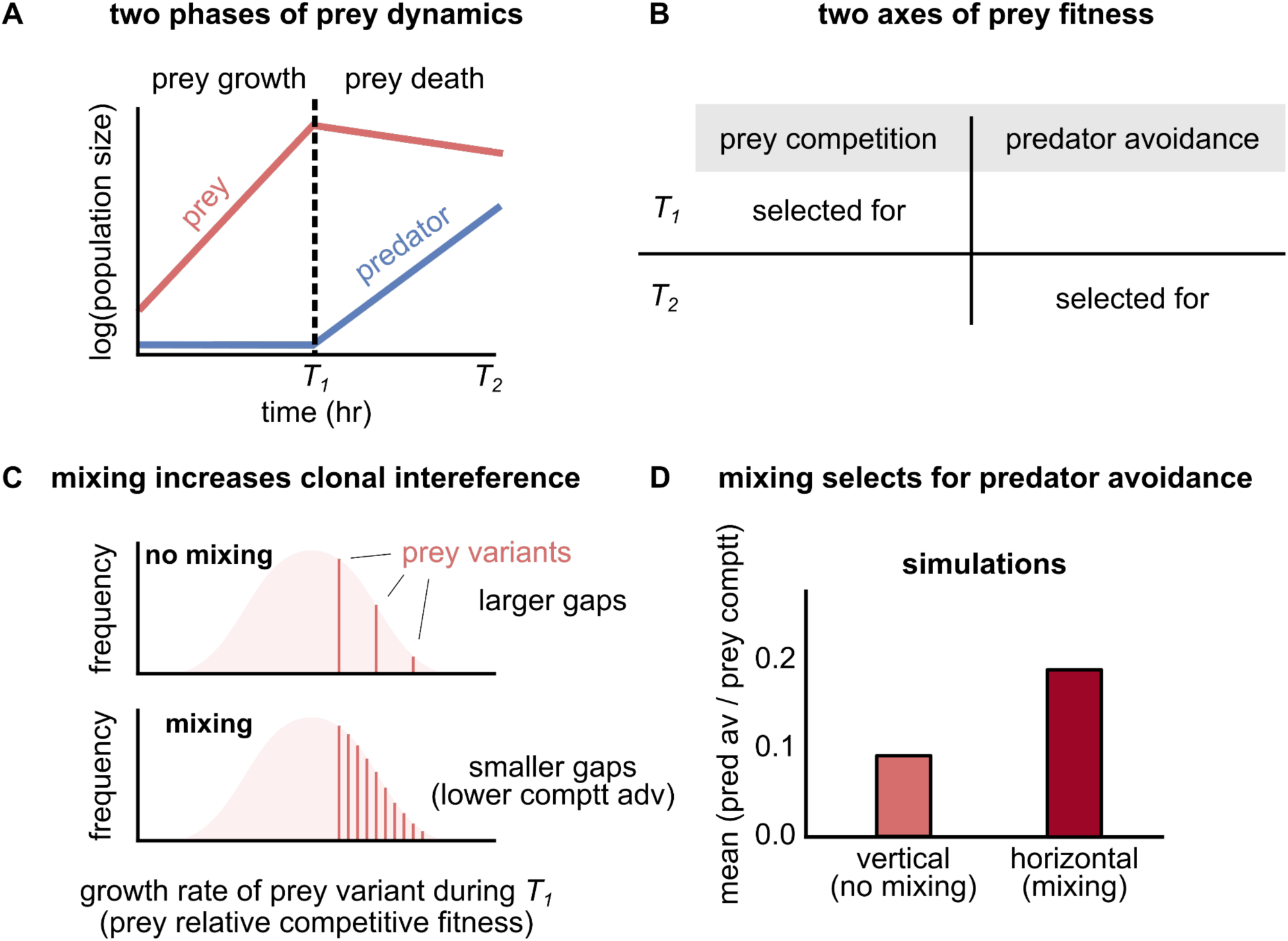
Mathematical model of population dynamics suggests that mixing increases clonal interference and preferentially selects for predator avoidance. A) Schematic illustrating the two dynamic phases of the prey and predator in each growth cycle. The first phase, lasting time *T1*, exclusively involves exponential prey growth (linear on a log axis), while the second phase, lasting time *T2*, involves predator growth and consequent prey death (Methods). B) In each dynamic phase, different aspects of the prey affect its fitness. In the prey growth phase, prey competition dominates, while during prey death, predator avoidance dominates. C) Schematic illustrating how mixing increases clonal interference and slows the evolution of prey competition. Shown are hypothetical growth rate distributions for prey subpopulations. Vertical lines show hypothetical prey variants. Mixing increases the diversity of competing prey variants, lowering the gaps between their growth rates and lowering the relative competitive advantage of prey variants with large growth rates. Thus, prey variants with decreased death rates have a larger selective advantage. D) Results from simulations of our model (for details see Methods) suggest that mixing selects for variants with increased predator avoidance over prey competition, providing an explanation for our observations.

The relative fitness of prey in each of the two phases operated along two distinct measures of fitness in our model. During prey growth, prey fitness was determined primarily by prey competition, i.e., the growth rate of each prey subpopulation relative to the others (Figure 4B). During predator growth, prey fitness was set primarily by predator avoidance, i.e., the death rate of each prey subpopulation (Fig. 4B and Methods for details of the model). Thus, the fitness for each prey subpopulation was determined by two parameters. Our model simulated population dynamics over these growth cycles for a population comprising several prey subpopulations whose prey competition (growth rate) and predator avoidance (death rate) were randomly picked from an appropriate distribution (details in Methods). To simulate mixing (horizontal) and no mixing (vertical) regimens, we mimicked our experimental protocol (Methods).

Simulations of our model reproduced our experimental observations: mixed populations showed greater predator avoidance and lower relative prey competitive ability (Figure 4C and 4D). Further, the model explained why mixing suppressed prey competition. The key idea is that mixing increases the number of competing prey subpopulations, which increases clonal interference (Figure 4D). This slows down the takeover of prey with a growth advantage and favors prey with improved predator avoidance. Thus, our model suggests that mixing increases clonal interference among competing prey, potentially explaining the different experimental outcomes in vertical and horizontal regimens.

### Prey populations from vertical and horizontal treatment show distinct evolutionary dynamics

Together, empirical observations and theoretical predations suggest that the prey populations in the vertical transfer regimen are under fluctuating selection. This was because during each growth cycle interference competition against conspecifics was under selection during the first growth phase and resistance to predation was under selection in the second growth phase (Figure 4A). This resulted in the emergence of distinct genotypes with different degrees of resistance to *M. xanthus* predation in vertical treatment. Since the benefits of better intra-prey population competitive ability during the growth phase were limited in the horizontal treatment, prey populations from the horizontal regimen were probably under directional selection, which selects for lesser phenotypic diversity. If true, horizontally evolved prey populations should over evolutionary time keep on either increasing or maintaining high levels of resistance against *M. xanthus*. While vertically co-evolved prey populations are likely to not show a trend of increasing resistance to predation over evolutionary time.

To test this hypothesis, we isolated *M. xanthus* populations after twenty rounds of coevolution. These *M. xanthus* populations were then used to test the resistance of evolved *E. coli* populations from three distinct evolutionary time points (i.e., after ten, sixteen and twentieth transfer). In line with our hypothesis, we observed that after sixteen transfers, populations of horizontally coevolved prey were on average 1.06 log-fold more resistant to predators than the ones from the tenth transfer. Further, though not statistically significant, differences were observed between prey at the sixteenth and twentieth transfer, prey at the sixteenth transfer on average seemed more sensitive to predation than the ones after twenty transfers (Figure 5A, ANOVA and Tukey’s post-hoc). Whereas the vertically coevolved prey did not show differences in the levels of resistance to predation from transfer-ten to transfer-twenty (Figure 5A, ANOVA and Tukey’s post-hoc). These data, coupled with our observations that within-population variation among clones from the horizontally coevolved prey populations is less (Figure 2), and that vertically evolved prey populations are internally heterogeneous (i.e., show diversity in predation resistance phenotypes) (Figure 2), supports our hypothesis that the horizontally co-evolved prey populations tend to either continuously evolve higher level of resistance of maintain the levels of resistance over evolutionary time. Whereas vertically co-evolved prey populations evolve two strategies, i.e. evolution of superior intraspecies competitive fitness and resistance to predation. However, when *M. xanthus* population isolated from different evolutionary time points (i.e., tenth, sixteenth and twentieth) were cultured with *E. coli* population isolated after the twentieth transfer, no significant differences were observed in the predatory ability of the predators over evolutionary time from both the treatments. Here, the survival of terminally coevolved *E. coli* (CFU/mL) in the presence of predators from distinct evolutionary timepoints within respective treatments, was taken as a read-out of the predatory abilities of respective predator population (Figure 5B, ANOVA and Tukey’s post-hoc)

**Figure 5:**
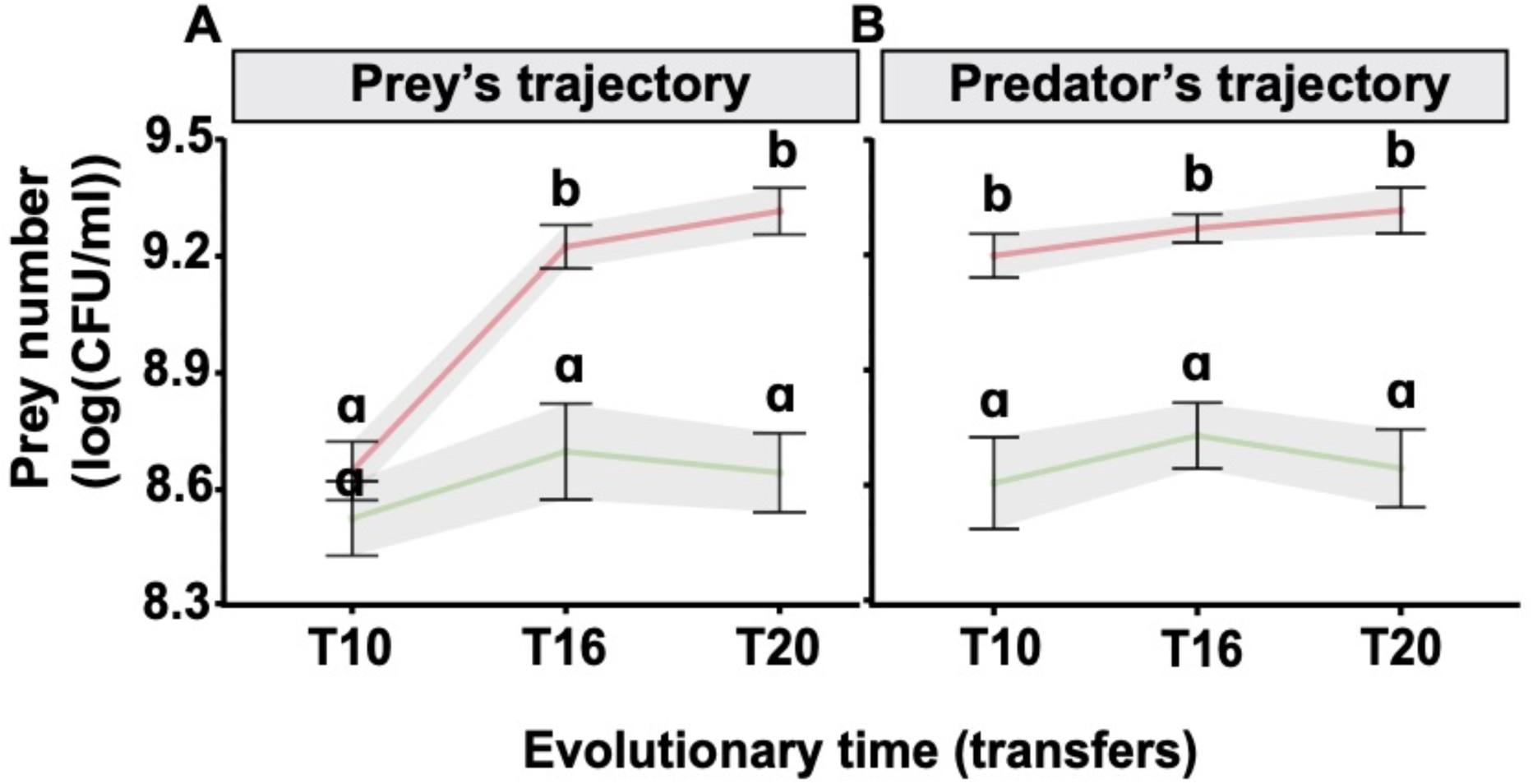
Prey from the horizontal selection regimen become increasingly more resistant over evolutionary time whereas the ones from the vertical regimen do not. A) Coevolved *E. coli* populations after ten (T10), sixteen (T16), and twenty (T20) transfers from vertical (green line) and horizontal (red line) regimens were analysed for changes in the resistance to predation against terminally evolved (after twenty cycles) respective *M. xanthus* populations. B) Coevolved *M. xanthus* populations after ten (T10), sixteen (T16) and twenty (T20) transfers from vertical (green line) and horizontal (red line) regimens were analysed for the change in the predatory performance against terminally evolved (after twenty cycles) respective *E. coli* populations from either vertical or horizontal regimen. Data shown is the average log(CFU/ml) for prey survival against respective *M. xanthus* from both regimens. Error bars are 95% confidence interval, letters indicate significant differences within the regimen (ANOVA, Tukey’s post hoc test, n = 3).

### Switching the mixing regimen between the terminally evolved horizontal and vertical populations

Predictions from our model suggest that the distinct outcomes in the two selection regimens depend on the continuation of the selection regime. Thus, if mixing indeed resulted in directional selection in prey survival strategies, repeating the same evolution experiment again with the diversified populations from the vertical regimen should result in one or both of the following possibilities. First, reduced variation among clones after being propagated in the mixing treatment. And second, increased resistance to predation both at the population and isolate levels. To test these hypotheses, we propagated a terminally coevolved population (i.e., after twenty transfers) by Switching the transfer regimen - i.e., populations from the horizontal regimen were propagated vertically, and the vertically evolved populations were propagated in the horizontal regimen - (Supplementary figure S7). Similar to the experiments above, prey populations were isolated from the evolved prey-predator consortia (See methods) after they were propagated for sixteen transfers. On testing the ability to survive predation in the evolved preys against mixing, not only evolved a significantly higher ability to survive predation than their ancestors (Figure 6A, ANOVA and Tukey’s post-hoc) but also exhibited a lower within-population diversity than their ancestors (Figure 6B, ANOVA and Tukey’s post-hoc for difference between groups and also for differences within each of the groups). Moreover, consistent with our previous results, while the overall resistance to predation did not change much at a population level (Figure 6A, ANOVA and Tukey’s Post-hoc), in the prey population that was propagated under vertical regimen exhibited a greater degree of within-population variation than their ancestors (Figure 6B, ANOVA and Tukey’s post-hoc for difference between groups and also for differences within each of the groups). Together, these experiments reveal that irrespective of preexisting standing diversity, prey populations that experience repeated mixing will evolve to have reduced within-population diversity and higher resistance than the ones that evolved in the absence of mixing events.

**Figure 6:**
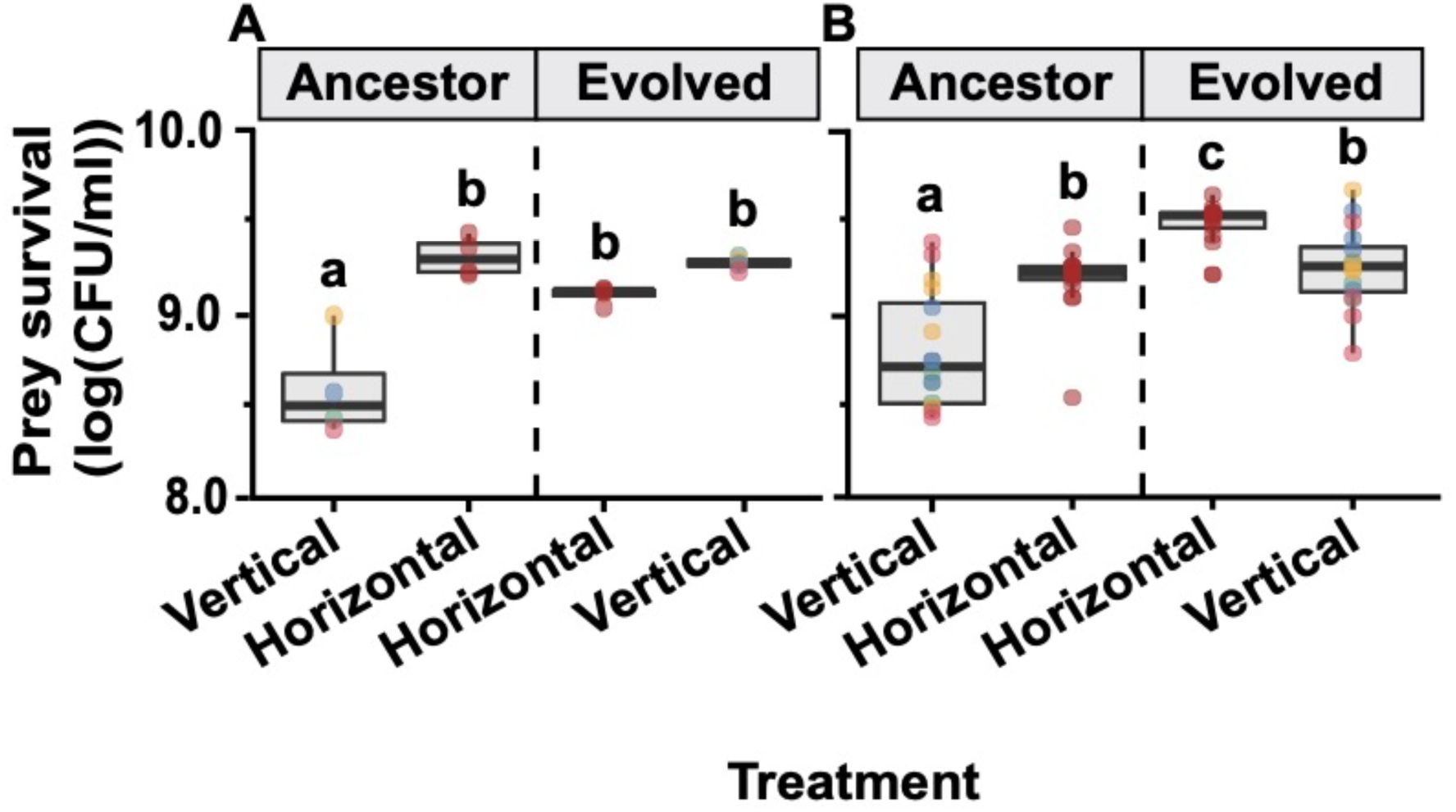
Mixing communities always results in evolution of higher resistance to predation while isolated communities evolve diversified survival strategies in prey population. A) Growth of ancestral vertically coevolved and horizontally coevolved *E. coli* population (T20 ancestors) along with the *E. coli* population after sixteen transfers in flipped regimen (Flipped: where vertically coevolved communities were further propagated under horizontal regimen and horizontally coevolved clones were further propagated under vertical treatment regimen), in presence of ancestral *M. xanthus* (ANOVA, Tukey’s post hoc test, n = 4). B) Growth of *E. coli* clones isolated from ancestral vertically coevolved and horizontally coevolved *E. coli* population (T20 ancestors) along with the *E. coli* clones isolated after sixteen transfers in flipped regimen (Flipped) in presence of ancestral *M. xanthus* (ANOVA, Tukey’s post hoc test, n = 4). Different color of dots in the vertical treatment represents distinct lineages.

## Discussion

Events that affect the mixing of communities are common in nature ^1,9,13,38–40^. However, their influence on the evolution of bacterial predator-prey interaction remains less explored ^1^. Here we used model predator-prey communities of bacteria to show that repeated mixing of predator-prey communities can result in the evolution of enhanced prey survival ability in the presence of the predator, relative to when communities are allowed to evolve in the absence of repeated mixing events. Surprisingly though, we could not detect increased predatory performance in the evolved populations of predators. Moreover, contrary to expectations we observed the evolution of reduced predatory performances in certain predator isolates. Hence the observations reported in our study do not follow the coevolutionary pattern of the evolutionary arms-race as expected from host-parasite, and host-pathogen interactions ^16,20,41^. Several factors such as social predation by *M. xanthus*, inter- vs intra-species competition, differences in the effective mutation supply in the two selection regimens, and asymmetry in the rate of evolution of prey and predator can explain the observed outcomes.

Two aspects of *M. xanthus* biology can introduce intra-species competition among predators. First, *M. xanthus* is a social predator that uses diffusible compounds to kill prey ^3,4,29,42^. Moreover, digestion of the prey is done extracellularly ^27,43^. Hence, both the killing factors, as well as the digested prey, are available as public goods to the individuals in the neighborhood. In our experiments, we find that most of the individual isolates from both treatments show similar predatory performance as their ancestors, except for a few clones, that perform worse than their ancestors. Hence, it seems possible that the apparent decrease in population productivity is mediated by the exploitation of public goods by the non-producers. If this is true, the use of non-social predators in a similar experimental setup should result in the evolution of increased predatory performance. Indeed, this is the case, predator-prey coevolution experiments using non-social *M. xanthus* isolates have resulted in enhanced predatory performance relative to the ancestral variants ^24^. Though not the focus of the current manuscript, it is an interesting opportunity to test the interaction dynamics between the coevolved efficient predator and coevolved less-efficient predator. Such population studies coupled with mechanistic understanding might reveal ecological and evolutionary mechanisms that lead to the evolution of scavenging strategies. That is the evolution of reduced prey-killing ability coupled with a strong dependence on the presence of efficient predators for better survival.

Microbial antagonism is pervasive in complex microbial communities ^44,45^. Hence, studies of such interactions have attracted tremendous interest in recent years to combat microbial infections and contaminations ^46–48^, and to better the understanding of microbial ecology ^24,32,49,50^. To our surprise, studies have used the terms host-parasite and predator-prey interactions interchangeably for identical interactions (for example bacteria and phage) ^38,51,52^. However, predator-prey interactions are distinct from host-parasite interactions in many ways ^51,54–58^ (for example: relationship intimacy, relationship duration, generation time, population size, and trophic level). Among them we feel two are relatively more significant than others. First, the population size of the exploiter and exploited is inverse for predator-prey interactions relative to that of the host-parasite interactions. Second, the generation time of the predator and prey is similar, whereas the generation time of parasites is significantly smaller than their hosts. Thus, the exploited partner has a faster rate of evolution in predator-prey interactions, which is the opposite for host-parasite interactions where the exploiter has a faster rate of evolution relative to the exploited partner. Here, we did not observe the evolution of increased predation efficiency among *M. xanthus* populations, whereas bacteria-phage (host-parasite) coevolution experiments have repeatedly shown the evolution of increased virulence ^1,20,53^. The differences we find here relative to the ones reported in the previous studies ^1,19,52^ in which the exploiter (parasite) evolves faster might be attributed to the differences in the relative rate of evolution of exploiter to the exploited partner.

The effect of population mixing has been studied in multiple instances ^1,59^. Whereas the effect of mixing of predator-prey communities has not been studied in as much detail. Interestingly, we observed that the evolution of prey bacteria followed a similar fate to previous reports in which the effect of population mixing was studied on the evolution of bacterial host-parasite interaction ^1^. Albeit with one key difference, i.e., unlike in the host-parasite evolution experiment in which vertical propagation either selected for resistance to phage in some replicate lineages or intraspecies competitive fitness in others, we observed that both of these strategies simultaneously evolved within each of the replicate population. Further, we observed that repeated mixing resulted in directional selection on the evolution of resistance against the exploiting partner. And unmixed communities, like in previous report ^24,32^, selected for the coexistence of distinct survival strategies that include the evolution of resistance and sensitivity to the exploiter, which is suggestive of fluctuating selection. If these trends observed in our experiments are indeed solely a result of the repeated mixing of communities, it is expected that when the diversified vertically coevolved communities are propagated under horizontal treatment regimen, homogenization of the survival strategies should be observed conferring overall increased resistance to predation. When the transfer regimen was flipped from horizontal to vertical, diverse survival strategies should emerge from a more homogeneously distributed population. Consistent with our expectation we observe an increased resistance to predation in our flipped horizontal treatment and diversification in our vertical treatments. Thus, supporting our previous observation and further establishing that mixing predominantly will result in an increased resistance to predation in prey irrespective of whether the ancestral population is isogenic or has a standing diversity.

Though the experimental approaches to studying the effect of population mixing are numerous, subtle differences between them introduce differential importance to the spatial structure. For example, population mixing within a single spatially structured arena will allow the mixing of distinct genotypes but will not affect the overall availability of diffused shared resources such as public-good ^60–62^. Whereas mixing of populations as done in this study will result in the mixing of distinct lineages repeatedly relative to the vertically propagated communities in which identical genotypes interact with each other over evolutionary time. Separation of populations during the growth phase and mixing at each transfer will result in increased mutational supply even after segregation of the populations. Thus, the design of the evolution experiment allows the horizontal population to behave as a single large metapopulation. Thus, we used a mathematical modelling approach to demonstrate that the mixing events, as expected, will effectively allow the horizontal populations (using the growth phase) to sample a larger number of mutations. The model further shows that this results in changes in the dynamics of interference competition. These changes in the dynamics of interference competition resulted in the selection for resistance to predation. This was because of the diminishing adaptive benefits of resource competition. The model further predicted that the outcomes of the evolution experiment depend on the continuation of regimens. That is, if the populations evolved in mixed treatment are propagated as unmixed, then it is expected that the populations will evolve diverse adaptive strategies like in the vertical treatment. We demonstrate that this outcome is indeed true by performing additional evolution experiments in which the selection regimens were switched.

The logistics of our evolution experiment involved aspects of evolution in the spatially structured as well as unstructured environment. For example: limited diffusion of resources is an important distinguisher between structured and unstructured environments, which has been shown to strongly affect the evolution of synergistic ^25,63^ as well as antagonistic interactions ^64^. The transfer protocol used in this study involved population mixing even in the vertically transmitted regimen, thus limiting the evolution of interaction networks. However, the probability of the evolution of synergistic interactions was most likely higher in the vertically propagated regimen than in the horizontal regimen. This is because of the mixing of multiple lineages before each transfer in the latter.

Interestingly, one of the four vertically propagated coevolving prey populations also demonstrated high levels of resistance to predation, which was comparable to horizontally propagated populations. Thus, vertically propagated prey populations might explore distinct evolutionary fates that range from the evolution of higher levels of resistance to *M. xanthus*, to the evolution of within-population diversity with both resistant and sensitive individuals. This finding is not surprising as similar effects of differential effects of bacterial predators, protists, and phages on bacteria have been demonstrated previously ^32,65–67^. Nonetheless, unlike prey populations that evolved in horizontal regimen, the vertical population with high levels of resistance to predation harboured intraspecies diversity (prey with low and high resistance to predation) which was similar to the other vertically propagated coevolving *E. coli* populations. Together, it seems that the evolution of the homogenous predation-resistant-prey population in horizontally propagated lines indicates that the predation-resistant individuals will outcompete predation-sensitive ones when communities are repeatedly mixed.

In recent years the call for harnessing the potential of microbial predators to control microbial contaminations and infections has generated tremendous interest. Thus, it is important to understand the effect of microbial predation on the evolution of prey species. In this context, it is unavoidable that distinct predator-prey populations might experience mixing events which might change the effective supply of mutations. Here we demonstrate that because of mixing events microbial predator-prey interactions might have a strong influence on the evolutionary outcomes for prey populations and that mixing will most likely have a stronger influence on the evolution of prey bacteria but not the predator.

## Supporting information

E coli Genomics

M. xanthus genomics

## Acknowledgements

The authors thank Deepa Agashe and Gregory Velicer for providing *E. coli* (MG1655) and *M. xanthus* (GJV1) strains, members of the Bacterial Ecology and Evolution lab at the Indian Institute of Science for helpful discussion. We also thank DST-FIST for support to the Department of Microbiology and Cell Biology, UGC Centre for advanced study, and the DBT-IISc partnership. AG acknowledges support from the Ashok and Gita Vaish Junior Researcher Award, as well as the Government of India’s DBT Ramalingaswami Fellowship. This study was mainly supported by the funds from India Alliance Intermediate fellowship to SP (IA/I/20/1/504921) and HGK-IYBA grant (BT/13/IYBA/2020/15) to SP.

## Author contribution

SP conceived the project. SS, BB, and SP designed experiments. SS, BB, and JL performed experiments. SS, BB and JL analyzed the data. AG performed the mathematical modelling. SS wrote the first draft of the manuscript. SP edited the manuscript.

## Methods

### Semantics

We define predation-prey interaction as an antagonist interaction in which the exploiting partner kills the exploited partner to acquire nutrients. Predator-prey interactions are distinct from host-parasite interactions in many ways ^51,54–58^ (for example: relationship intimacy, relationship duration, generation time, population size, and trophic level). Importantly, we argue that two factors most likely to differentiate the evolutionary dynamics of host-parasite interactions relative to that of predator-prey interactions are.

1. The overall population size of the predator is similar or lower relative to the prey. Thus, prey is generally present in higher numbers than predators.
2. Unlike host-parasite interactions in which parasites have smaller generation time, predator and prey organisms generally have similar generation times.

Thus, unlike host-parasite or host-pathogen interactions, the exploited organisms evolve faster than the exploiters in predator-prey communities.

### Strains used

*M. xanthus* (GJV1) and *E. coli* (MG1655) were used as predator and prey respectively. *M. xanthus* strain used in this study was provided by Dr. Gregory Velicer’s laboratory at ETH Zurich, Zurich, Switzerland. Whereas *E. coli* (MG1655) strain was provided by Dr. Deepa Agashe’s laboratory at the National Centre for Biological Sciences, Bangalore, India.

### Culture condition

All plates were incubated at 32 °C, all liquid cultures were incubated at 32 °C, 200 RPM. *M. xanthus* strains were stored as frozen glycerol (20%) stocks in CTT ^68^ medium, and *E. coli* strains were stored in frozen glycerol (20%) stocks in LB medium (HIMedia Laboratories Pvt. Ltd, India).

To initiate the evolution experiment *M. xanthus* glycerol stock was inoculated on CTT medium with 1.5 % agar, incubated for 3-5 days, inoculated from colony edge into 8 mL liquid CTT medium, and incubated for 24 h at 200 RPM. Grown liquid culture of *M. xanthus* was dilution plated on CTT 0.5 % agar plates and colonies (subclones) were isolated, stored as freezer stock and distinct subclones were used as biological replicates to initiate the evolution experiment.

Similarly, *E. coli* strains were inoculated on LB medium with 1.5 % agar, incubated for 1-day, individual colonies were inoculated into 8 mL liquid LB medium, and incubated for 24 h at 200 RPM. Each colony (subclone) of *E. coli* served as an ancestor for an independent replicate line.

For experiments performed to analyse evolved population/clones of *M. xanthus*, glycerol stock of *M. xanthus* was spotted on CTT medium with 1.5% agar plates and incubated at 32°C for 3-4 days. Once the swarm appeared, cells from the outer edge (actively growing and swarming) were inoculated in CTT liquid and incubated under shaking conditions at 32 °C for 20 h. For *E. coli*, cultures were grown in M9 (MP Biomedicals, United States) + 0.2% glucose (ThermoFisher Scientific, India) liquid, prior to setting all experiments.

### Evolution experiment 1

Protocol for the evolution experiment is demonstrated in Figure S1. Evolution experiment was performed in 50 mL flasks containing 10 mL M9 medium + 1.5 % agar medium (M9 salts (MP Biomedicals) + 0.2 % glucose + 0.01mM CaCl_2_ (ThermoFisher Scientific, India) + 0.2mM MgSO_4_ + 1.5% bacto agar (DIFCO, BD Biosciences, India)). Glucose in the M9 medium supports the growth of *E. coli* whereas *M. xanthus* cannot utilise glucose as a carbon source, and hence depend on their predatory ability to feed on the growing population of *E. coli*.

To initiate the evolution experiment, the densities of *E. coli* and *M. xanthus* were adjusted to ∼10^6^ cells/mL and ∼2*10^10^ cells/mL respectively by resuspending in M9 buffer (M9 salts (MP Biomedicals) + 0.01mM CaCl_2_ + 0.2mM MgSO_4_). 50 µL culture of prey and predator was spread across the agar bed with 9-12 sterile glass beads. For the *E. coli* only control 50 µL of *E. coli* culture was spread on the agar beds along with 50 µL of M9 buffer (M9 salts (MP Biomedicals, United States) + 0.01mM CaCl_2_ + 0.2mM MgSO_4_). For predator-only control, agar beds were supplemented with 5% dead *E. coli.* Dead *E. coli* were obtained by autoclaving 10 mL of 1.0 OD culture of *E. coli* for 20 minutes. All agar beds were prepared 1 day prior to the transfers. After 96 h of incubation at 32°C the predator-prey communities were harvested by adding 4 mL of M9 buffer followed by shaking for ∼30 mins at 200 RPM to resuspend the growing culture. For the vertical transfer regimen, 200 μL (5 %) of the resuspended culture from each flask was transferred to a fresh flask and spread using glass beads. In horizontal transfer regimen, all four replicate cultures of the respective treatment were pooled in a single 15 mL conical centrifuge tube, mixed, 200 μL of these pooled cultures were redistributed in each of the four replicate flasks with fresh medium and spread with the help of 9-12 glass-beads. The evolution experiment was carried out for 20 such transfer cycles with populations being stored at regular intervals in 20 % glycerol stocks at -80 °C.

### Isolation of clones from evolved populations of *M. xanthus* and *E. coli*

To isolate the evolved population of *E. coli*, a fraction of frozen glycerol stock was inoculated in 5 mL LB liquid medium (with 1 % NaCl instead of 0.5 %), incubated at 32 °C at 200 RPM for 7-8 h, and frozen as 20 % glycerol stock. *M. xanthus* cells are sensitive to the presence of salt and hence do not grow in these conditions, whereas cells of *E. coli* do.

To isolate *M. xanthus* population, frozen glycerol stock was inoculated in 8 mL CTT liquid with Gentamicin (Sisco Research Laboratories Pvt. Ltd, India) (10 μg/mL), incubated at 32 °C at 200 RPM for 18-20 h, and frozen as 20 % glycerol stock. Being naturally resistant to Gentamicin, *M. xanthus* can grow in the presence of Gentamicin.

Next, to isolate *E. coli* clones glycerol stocks of frozen *E. coli* populations were dilution plated on LB + 0.5 % agar (Agar agar, ThermoFisher Scientific, India), incubated at 32 °C for 18 h. Distinct colonies were isolated as clones, inoculated in liquid LB medium, incubated at 32 °C at 200 RPM for 7-8 h, and stored as glycerol (20 %) stocks at -80 °C.

To isolate *M. xanthus* clones glycerol stocks of frozen *M. xanthus* populations were pour-platted in CTT + 0.5 % agar, and incubated at 32 °C for 96 h. Distinct colonies were isolated as subclones, inoculated in liquid CTT medium, incubated at 32 °C at 200 RPM for 20 h, and stored at glycerol (20 %) stocks at -80 °C.

### Predation assay

Predatory performance can be measured as the growth of M. *xanthus* when prey is the only source of nutrient, or growth of *E. coli* in the presence of *M. xanthus* relative to its absence. In our study, we used growth of *E. coli* as a measure of predation by *M. xanthus*. This is because *M. xanthus* is a sticky biofilm forming bacterium. Therefore, estimates of Colony Forming Units (CFU) are generally unreliable, and a single colony might emerge from a group of cells resulting in significant underestimation of *M. xanthus* cell numbers.

All predation assays were performed on 10 mL M9 + 1.5 % agar beds (M9 salts (MP Biomedicals) + 0.2 % glucose + 0.01 mM CaCl_2_ + 0.2 mM MgSO_4_ +1.5 % Bacto agar) in 50 mL conical flasks with 9-12 sterile glass beads. Media was poured into the flask one day prior to the experiment. All *E. coli* cultures were grown to log phase in liquid M9 media (M9 salts (MP Biomedicals) + 0.2 % glucose + 0.01 mM CaCl_2_ + 0.2 mM MgSO_4_) prior to setting up the experiments. The *M. xanthus* cultures were grown to log phase in liquid CTT media. The prey cell density was adjusted to 0.1 OD (∼10^6^ cells/mL) and predator cell density was adjusted to 5*10^9^ cells/mL in TPM buffer (10 mM Tris pH 8.0, 8 mM MgSO_4_, 1 mM K_2_HPO_4_ – KH_2_PO_4_), 50μL of these cultures were then inoculated on the M9+1.5 % agar beds and spread with sterile glass beads. For monoculture of *E. coli,* 50 μL of cell density adjusted culture was inoculated along with 50 μL TPM buffer and spread on the M9+ 1.5 % agar medium. The cultures were incubated for 96 h at 32 °C. Following the incubation period, cells were harvested by adding 4 mL of TPM buffer with 9-12 sterile glass beads followed by ∼30 min shaking at 200 RPM. Viable number of *E. coli* was determined by dilution plating the resuspended cell suspension in LB + 0.5 % agar.

### Competition assay

#### Competition between ancestors with and without Kanamycin marker

*E. coli* ancestral strains with and without Kanamycin marker were grown to log phase in LB liquid medium and 1:1 ratio of 0.1 OD adjusted culture of each strain was co-inoculated to initiate the experiment. The competition experiment was performed on 10 mL M9 + 1.5 % agar beds (M9 salts (MP Biomedicals) + 0.2 % glucose + 0.01 mM CaCl_2_ + 0.2 mM MgSO_4_ + 1.5 % Bacto agar) in 50 mL conical flasks with 9-12 sterile glass beads. The culture was spread evenly on the agar-beds using glass beads and incubated for 24 h at 32 °C. Post incubation, the cells were harvested by adding 4 mL TPM buffer with ∼30 min shaking at 200 RPM. Viable counts of Kanamycin sensitive and resistant *E. coli* were determined by dilution plating on LB + 0.5 % agar and LB + 0.5 % agar with Kanamycin (40 μg / mL) (Merck Germany).

#### Competition between evolved clones and Kanamycin marked ancestors

Four *E. coli* clones were isolated from each of the vertically evolved population, sixteen from each from horizontally co-evolved and mono-evolved populations. Each of these clones were competed against their respective ancestors, in the presence or absence of ancestral *M. xanthus.* For these experiments we used Kanamycin-resistant ancestor *E. coli*, which allowed us to distinguish them from the evolved clones. Competition experiments were performed on 10 mL M9 + 1.5 % agar beds (M9 salts (MP Biomedicals) + 0.2 % glucose + 0.01 mM CaCl_2_ + 0.2 mM MgSO_4_ + 1.5 % Bacto agar) in 50 mL conical flasks with 9-12 sterile glass beads. All *E. coli* cultures were grown to log phase in M9 liquid media, 25 μL of 0.1 OD (∼10^6^ cells/mL) adjusted culture of each evolved and ancestor *E. coli* were co-inoculated, either with (50 μL of 5*10^9^ cells/mL) or without (50 μL TPM buffer) ancestral *M. xanthus*, spread on the agar medium with sterile glass beads. These were incubated for 96 h at 32 °C and harvested by adding in 4 mL TPM buffer with ∼30 min shaking at 200 RPM. Viable counts of ancestor and evolved *E. coli* were determined by dilution plating on LB + 0.5 % agar and LB+0.5% agar with Kanamycin (40 μg/mL) (Merck Germany).

#### Determination of prey-predator growth dynamics in co-cultures experiments

To demonstrate the relative growth dynamics of the prey and predator when co-cultured together in media containing only glucose as a carbon source, the prey numbers and predator densities were tracked for 4 days. Since only *E. coli* can utilise glucose and *M. xanthus* cannot. The only available source of nutrition for *M. xanthus* was the prey population. Additionally, given the differences in generation times of *E. coli* and *M. xanthus*, their growth in the coculture is expected to be temporally segregated. Hence, it is expected that *E. coli* will have distinct growth and death phases. Thus, to determine the changes in the population size of prey bacteria during the four-day transfer cycle, prey counts were obtained every 24 h from 0 to 96 h. To do so, 50 μL of 0.1 OD (∼10^6^ cells/mL) adjusted culture of ancestor *E. coli* was co-inoculated, either with (50 μL of 5*10^9^ cells/mL) or without (50 μL TPM buffer) ancestral *M. xanthus*, and spread on 10 mL of M9 + 1.5 % agar beds (M9 salts (MP Biomedicals) + 0.2 % glucose + 0.01 mM CaCl_2_ + 0.2 mM MgSO_4_ + 1.5 % Bacto agar). The cultures were spread on the agar medium with the help of sterile glass beads, incubated at 32 °C, and harvested by adding in 4 mL TPM buffer with ∼30 min shaking at 200 RPM. Viable counts of *E. coli* were determined by dilution plating on LB + 0.5 % agar. Because of the destructive sampling approach, distinct prey-predator microcosms were set up for each time point.

Since exopolysaccharides secreted by *M. xanthus* result in the formation of biofilms that are difficult to disrupt, it reduces the reliability of the obtained *M. xanthus* viable counts, using similar methods as for *E. coli*. Therefore, we used the swarm sizes as a proxy to determine the relative densities of *M. xanthus* in the cocultures. Swarming of *M. xanthus* on hard agar surfaces is known to be a positive density-dependent trait. Hence, it is expected that as the relative numbers increase, the swarm sizes will also increase. To estimate the swarm sizes, post harvesting of the cocultures in TPM buffer (as described above), 10 μL of the culture was spotted on CTT hard agar (1.5 Bacto agar) surface in 90 mm plates. The initial swarm was marked after 24 h and the final swarms were marked after 120 h. The average distance between the initial and final swarms was recorded and presented as the swarm sizes.

#### Mathematical model of evolution in predator-prey communities

To explain our surprising experimental observation that variants with superior competitive fitness that did not show any increase in their ability to resist predation evolved only in the vertical treatment and not in the horizontal treatment, we developed a mathematical model of population dynamics. Our model followed the dynamics of prey subpopulations explicitly and the predator populations implicitly. Specifically, we assumed each growth cycle (which lasted time *T*) had two distinct periods: the first where prey subpopulations grew (lasting time *T_1_*) and the second where they died due to predation (lasting time *T_2_*). In each period, each subpopulation (labeled α) had an exponential growth rate *g_α_* and death rate *d_α_* respectively, both drawn randomly and independently from a normal distribution with means 0.7 hr^-1^ and 0.2 hr^-1^ respectively, and standard deviations 0.3 hr^-1^ and 0.2 hr^-1^ respectively (disallowing negative rates). With these the growth of each subpopulation was described by the following equation:

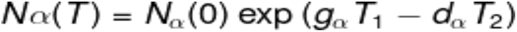

Inspired by evolutionary ecology, our model implemented evolution by introducing distinct “mutants” as prey subpopulations at the beginning of population dynamics and following their fate over time. We modeled 4 distinct communities, each seeded with 5 subpopulations of prey (chosen arbitrarily). At the beginning of each simulation, each subpopulation started with equal abundance 0.1 (arbitrary units) and grew according to the dynamics described above. After each growth cycle (time *T*), all subpopulations in each population were diluted by a factor *D = 20* (5% bottleneck) and grown again in a new cycle. In the vertical regimen, different communities were not mixed, while in the horizontal one, they were mixed and redistributed equally, mimicking our experiment. We continued these dynamics across multiple growth-dilution cycles till communities reached a steady state, after which we compared the mean ratio of the relative growth rates (quantifying prey competition) and death rates (quantifying predator avoidance) of each surviving subpopulation. We repeated and averaged our results over such 100 simulations with randomly chosen parameters and plotted the results (Fig. 4D).

#### Time-course assay

Predation assays (were performed with populations of coevolved prey and predator from both the transfer regimes (vertical and horizontal), respectively isolated from different evolutionary time. The populations of both prey and predator were isolated from the glycerol stocks stored at different transfers, namely, after transfer 10 (T10), transfer 16 (T16) and transfer 20 (T20). The prey, and the predator populations were isolated using a similar protocol as described earlier (See isolation of population and clones). To determine the change in prey survival over evolutionary time against the terminally coevolved predator, *E. coli* populations from different evolutionary times (T10, T16 and T20) were inoculated with the terminally evolved *M. xanthus* (after 20 transfers). The predation assay for each combination of prey and predator population was set up as discussed earlier (See predation assay). The change in prey survival over time was recorded for prey population by dilution plating on LB + 0.5 % agar. Similarly, the *M. xanthus* populations from all time points (T10, T16 and T20) were pitched against the terminally co-evolved *E. coli* population, to determine the changes in the predatory ability over evolutionary time. The survival of the terminally coevolved prey against *M. xanthus* from different evolutionary time-point was used as a measure of change in the resistance to predation over evolutionary time. Here, a lower prey survival implies better predatory ability, and a higher prey survival indicates worse predatory ability.

#### Flipped-regime evolution experiment (Evolution experiment 2)

Experimental protocol for the second evolution experiment is demonstrated in Supplementary figure S7. Evolution experiment was performed in 50 mL flasks containing 10 mL M9 + 1.5 % agar medium (M9 salts (MP Biomedicals) + 0.2 % glucose+ 0.01mM CaCl2 + 0.2mM MgSO4 + 1.5 % bacto agar). To initiate the second evolution experiment, glycerol stocks of the T20 population from evolution experiment 1 were thawed on ice and 5% of the culture was washed with M9 buffer two times and then resuspended in 200 μL M9 buffer. These cultures were then spread on fresh M9 + 1.5 % agar beds with the help of 9-12 sterile glass beads. Similar treatment was done for all populations from both mixed communities of prey and predator and monoculture prey (control) treatments. The mixed communities of prey and predator (coevolved) /prey only (mono-evolved) population were incubated in fresh media at 32 ^O^C for 96 h. This initial 96 h incubation was done to allow the stabilization of the communities after revival from freezer stocks. Following this both the mixed communities of prey-predator and prey only treatments were harvested in 4 mL of M9 buffer and shaking at ∼200 RPM for 30 mins to resuspend the growing cultures. 5 % of the resuspended culture was transferred to fresh M9 medium + 1.5 % agar medium under two different transfer regimens: Vertical transfer regimen and Horizontal transfer regimen (similar to the first experiment). However, in this experiment the transfer regimens of the evolved population from evolution experiment-1 were interchanged. The vertically coevolved/mono-evolved population were propagated under the horizontal transfer regimen, i.e., the population from replicate lines were pooled in a 15 mL centrifuge tube and then 5% of the resuspended culture was transferred to each of the 4 flasks with fresh media, at every transfer (ensuring a periodic mixing of population). Whereas the horizontally coevolved/mono-evolved population were propagated under the vertical transfer regimen, where suspended cultures from the 4 replicate lines were serially transferred to respective fresh M9 + 1.5 % agar medium. All the cultures were spread on the agar beds using the glass beads. The experiment was carried out for 20 such cycles with population being stored at regular intervals in the form of 20 % glycerol stocks in -80 °C. Evolved population of prey and predator was isolated after 16 transfer cycles using similar protocols as used for the isolation of population in the previous experiments.

### Genomic analysis

For the isolation of genomic DNA, *E. coli* and *M. xanthus* clones were grown in M9 + 0.2 % glucose and CTT liquid medium respectively at 32 °C, 200 RPM till they achieved optical density between 0.2 to 0.8 at 600 nm. These cultures were used for the extraction of genomic DNA using the QIAGEN genomic DNA isolation kit using 20/G column. Manufacturer’s instructions were followed for the isolation of genomic DNA. Next steps for sequencing were performed at Macrogen Inc (Korea). Before TrueSeq library preparation the quality control of the samples was done using Agilent TapeStation^R^ DNA Screen Tape (Agilent). The DNA samples were sequenced on HiSeq platform, for 150 bp paired-end sequencing. Ambiguous low-quality sequences were trimmed using Trimmomatic v0.39. The Genomic libraries were analysed using breseq^69^ against reference genome of *E. coli* MG1655 (GenBank Accession No. GCA_000005845.2) or *M. xanthus* DK1622 (GenBank Accession No.GCA_000012685.1) from NCBI genomic data bank. The breseq pipeline was used with all default parameters.

### Statistical analysis

All statistical analysis were performed in R (version 4.0.4). Levene’s test was performed with each set of data to check for the homogeneity of variance. For a p > 0.05 for Levene’s test, either an independent sample t-test or one sample t-test was performed wherever applicable, and the Benjamini-Hochberg method was used for correcting for multiple testing. Significant ANOVA in all analysis was followed by an SNK *post hoc* test. However, for a p < 0.05 for Levene’s test nonparametric tests were used for data analysis. The significant Kruskal-Wallis test was followed by Dunn’s *post hoc* test. The statistical test for the respective data set is mentioned in the figure legends.

