## Supplementry data for "Mixing selects for predation resistance in lab-evolved communities of bacterial prey and social predator *Myxococcus xanthus*"

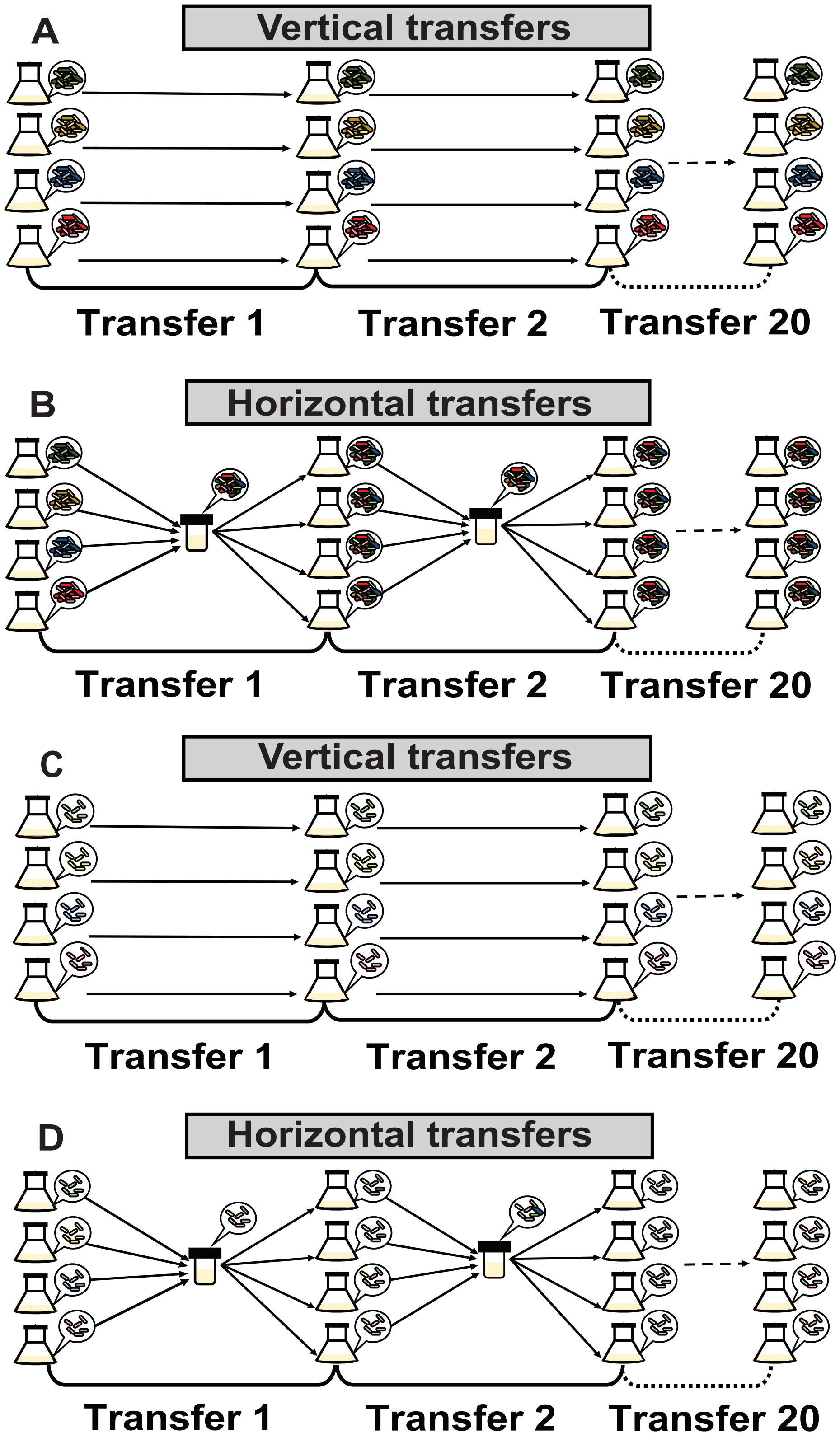
A
Vertical transfers
Transfer 1
Transfer 2
Transfer 20
B
Horizontal transfers
Transfer 1
Transfer 2
Transfer 20
C
Vertical transfers
Transfer 1
Transfer 2
Transfer 20
D
Horizontal transfers
Transfer 1
Transfer 2
Transfer 20

**Figure S1:** Four identical predator-prey consortia of *M. xanthus* and *E. coli* were serially propagated either horizontal or in vertical transfer regimen. (A) In vertical transfer regimen each of the four consortia were incubated for four days before being transfer to fresh medium without mixing. (B) Whereas in horizontal treatment all four communities were mixed after four days of incubation before redistributing them into four distinct consortia. Experiment was conducted for twenty transfers. Darker shades of colors represent *M. xanthus*, lighter shades represent *E. coli*, different colors represent identities of four distinct founding populations. (C and D) Similarly, as controls either the prey or predator was transferred under two regimens in absence of the other partner (Mono-evolved). Here the schematic only represents mono-evolved prey conditions (where preys evolved in absence of predator). We also had a similar treatment where predator was evolving alone (See methods).

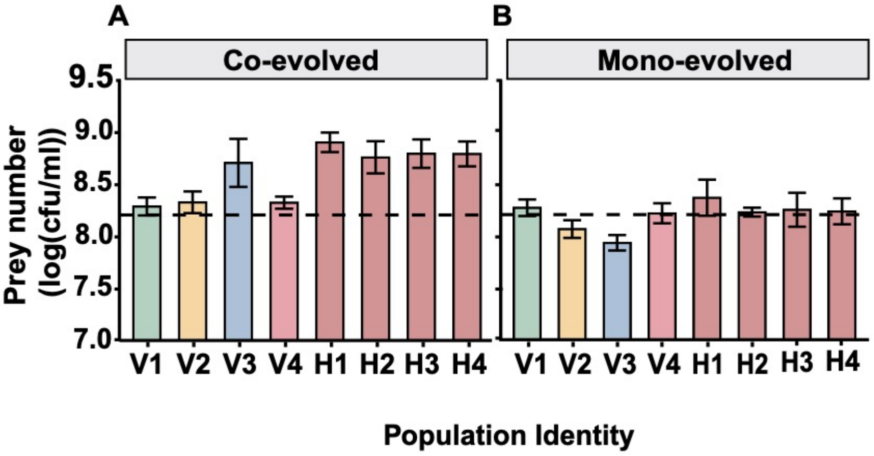


**Figure S2:** Growth of coevolved (A), and mono-evolved (B) populations of *E. coli* in the presence of *M. xanthus* were measured to assess the sensitivity of *E. coli* populations to *M. xanthus* predation. Green, yellow, blue and pink colors represent distinct vertical lineages (V1 to V4). Horizontal populations are shown in darker shade of pink (H1 to H4). Dashed line represents growth of ancestral strain of *E. coli* in the presence of *M. xanthus*. Error bars indicate 95 % confidence intervals (ANOVA between distinct populations, $p<0.05$, $n=4$).

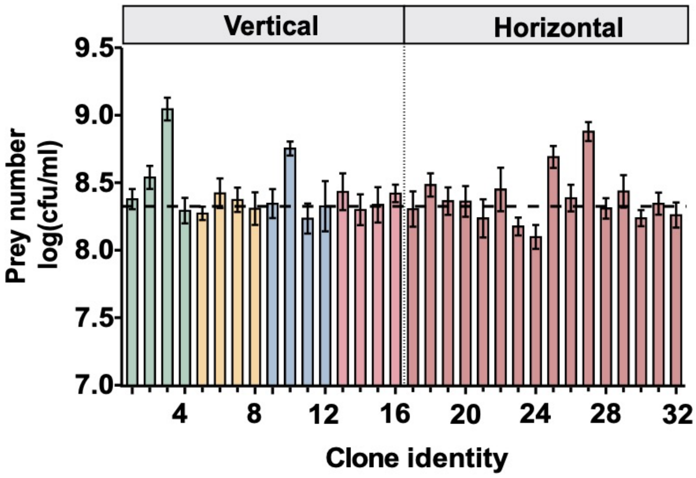


**Figure S3:** Growth of ancestral *E. coli* strain in the presence of distinct coevolved *M. xanthus* clones was measured as a measure predator performance of respective *M. xanthus* clone. Green, yellow, blue and pink indicate distinct lineages. Horizontal populations are shown in darker shade of pink. Dashed line represents growth of *E. coli* in the presence of ancestral *M. xanthus*. Error bars indicate 95 % confidence intervals. (See Supplementary Table T 1) (ANOVA and SNK post hoc test, n = 8).

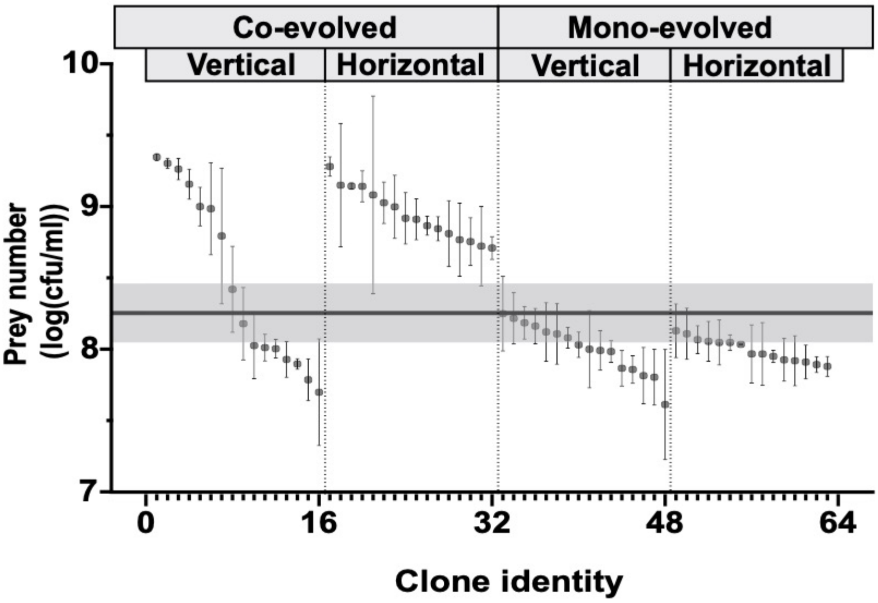


**Figure S4:** Growth of evolved and ancestral clones of *E. coli* in the presence of ancestral *M. xanthus* was quantified as a measure of their sensitivity to *M. xanthus* predation. Each dot represents mean growth of *E. coli*, error bars indicate 95 % confidence intervals, horizontal line at y= 8.36 and shaded region is mean survival of ancestral *E. coli* and associated 95 % confidence interval respectively n = 3. (ANOVA was done within clones from each treatment (See Supplementary table T2). Significant ANOVA was followed by SNK post-hoc test, n=3)

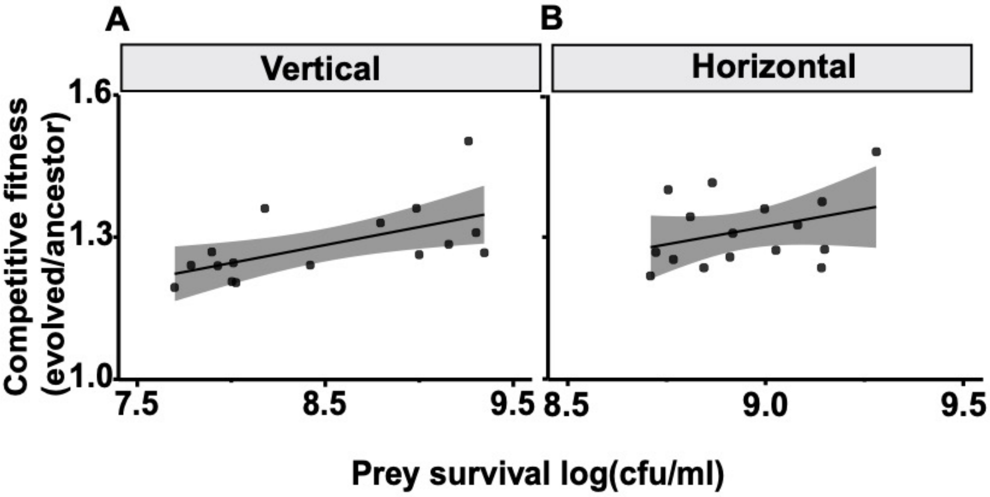


**Figure S5**: Growth of distinct *E. coli* isolates from vertically (A) and horizontally (B) coevolved populations in the presence of *M. xanthus* is plotted against their competitive fitness against ancestral *E. coli* in the presence of *M. xanthus*. Trend line is Pearson's correlation, and shaded region is 95 % confidence interval. R value for vertical transmission regimen is 0.6 ($p < 0.02$, and for the horizontal regimen R-value is 0.35 ($p < 0.2$), n = 3.

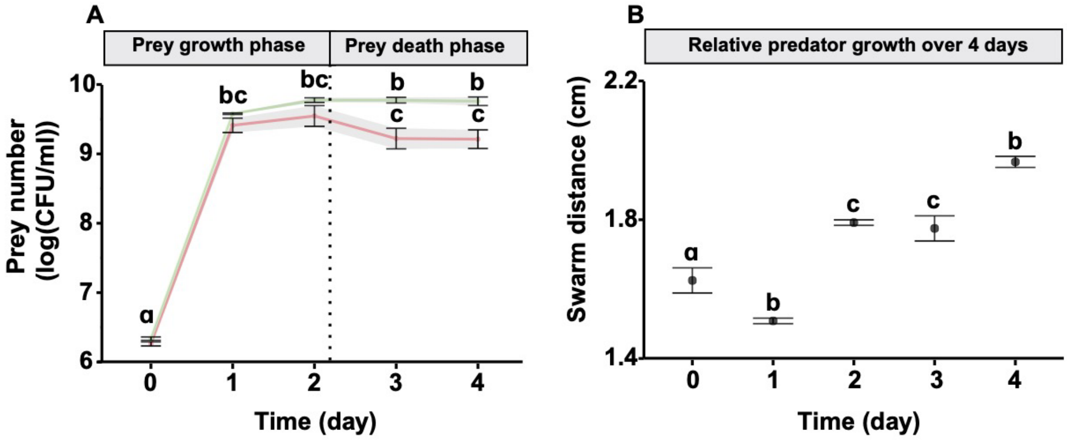


**Figure S6**: Prey -predator growth dynamics in co-culture experiments A) Growth of ancestral *E. coli* in the presence (red line) and absence (green line) of ancestral *M. xanthus* is plotted over a period of 4 days (different letters indicate differences between distinct datapoints, ANOVA for between different conditions across time points followed by SNK post hoc test, n = 3). B) Growth of *M. xanthus* represented as swarm sizes over 4 days when cocultured with *E. coli* (ANOVA between the growth at different time points followed by SNK post hoc test, n = 3).

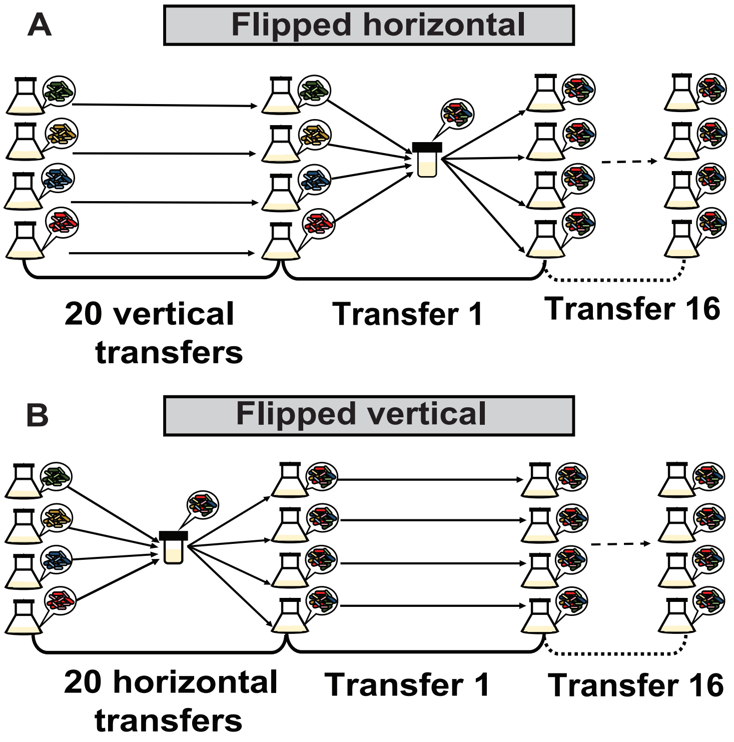


**Figure S7:** Terminally coevolved (after twenty transfers) communities from evolution experiment 1 (Supplementary figure S1) was further propagated by exchanging their transfer regimen. Where, vertically coevolved prey-predator consortia were now propagated under the horizontal transfer regimen (allowing repeated mixing of these consortia). Whereas horizontally coevolved prey was propagated under the vertical transfer regimen. These communities were propagated for sixteen transfers.

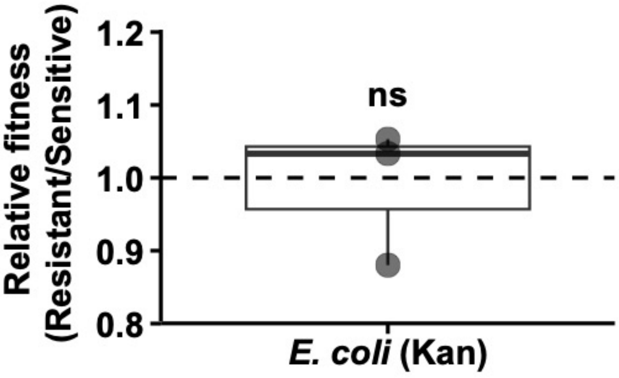


**Figure S8:** Competitive fitness of the Kanamycin E. coli strain relative to the unmarked ancestral *E.* coli strain when competed in 1:1 ratio (One- sample t-test, $p^{[FDR\ corrected]} > 0.8$, n =3).

| Clone Identity | Treatment | Group |
|---|---|---|
| V1 | Vertical | bcd |
| V2 | Vertical | bcd |
| V3 | Vertical | a |
| V4 | Vertical | cd |
| V5 | Vertical | cd |
| V6 | Vertical | bcd |
| V7 | Vertical | bcd |
| V8 | Vertical | cd |
| V9 | Vertical | bcd |
| V10 | Vertical | abc |
| V11 | Vertical | cd |
| V12 | Vertical | cd |
| V13 | Vertical | bcd |
| V14 | Vertical | cd |
| V15 | Vertical | bcd |
| V16 | Vertical | bcd |
| H1 | Horizontal | cd |
| H2 | Horizontal | bcd |
| H3 | Horizontal | bcd |
| H4 | Horizontal | bcd |
| H5 | Horizontal | cd |
| H6 | Horizontal | bcd |
| H7 | Horizontal | cd |
| H8 | Horizontal | cd |
| H9 | Horizontal | abc |
| H10 | Horizontal | bcd |
| H11 | Horizontal | ab |
| H12 | Horizontal | cd |
| H13 | Horizontal | bcd |
| H14 | Horizontal | cd |
| H15 | Horizontal | bcd |
| H16 | Horizontal | cd |

**Supplementary Table T1:** Shows the statistically significant differences in the survival of ancestor *E. coli* when cocultured with coevolved *M. xanthus* clones from both the treatments. Letters indicate significant differences.

| Clone identity | Treatment | Condition | ANOVA(p-value) | Group |
|---|---|---|---|---|
| V1 | Vertical | Coevolved | 2.03e-06 | bcd |
| V2 | Vertical | Coevolved | | cd |
| V3 | Vertical | Coevolved | | d |
| V4 | Vertical | Coevolved | | abc |
| V5 | Vertical | Coevolved | | abcd |
| V6 | Vertical | Coevolved | | a |
| V7 | Vertical | Coevolved | | cd |
| V8 | Vertical | Coevolved | | a |
| V9 | Vertical | Coevolved | | cd |
| V10 | Vertical | Coevolved | | d |
| V11 | Vertical | Coevolved | | cd |
| V12 | Vertical | Coevolved | | abc |
| V13 | Vertical | Coevolved | | ab |
| V14 | Vertical | Coevolved | | abcd |
| V15 | Vertical | Coevolved | | cd |
| V16 | Vertical | Coevolved | | a |
| H1 | Horizontal | Coevolved | 0.957 | ns |
| H2 | Horizontal | Coevolved | | |
| H3 | Horizontal | Coevolved | | |
| H4 | Horizontal | Coevolved | | |
| H5 | Horizontal | Coevolved | | |
| H6 | Horizontal | Coevolved | | |
| H7 | Horizontal | Coevolved | | |
| H8 | Horizontal | Coevolved | | |
| H9 | Horizontal | Coevolved | | |
| H10 | Horizontal | Coevolved | | |
| H11 | Horizontal | Coevolved | | |
| H12 | Horizontal | Coevolved | | |
| H13 | Horizontal | Coevolved | | |
| H14 | Horizontal | Coevolved | | |
| H15 | Horizontal | Coevolved | | |
| H16 | Horizontal | Coevolved | | |
| V1 | Vertical | Mono-evolved | 0.695 | ns |
| V2 | Vertical | Mono-evolved | | |
| V3 | Vertical | Mono-evolved | | |
| V4 | Vertical | Mono-evolved | | |
| V5 | Vertical | Mono-evolved | | |
| V6 | Vertical | Mono-evolved | | |
| V7 | Vertical | Mono-evolved | | |
| V8 | Vertical | Mono-evolved | | |
| V9 | Vertical | Mono-evolved | | |
| V10 | Vertical | Mono-evolved | | |
| V11 | Vertical | Mono-evolved | | |
| V12 | Vertical | Mono-evolved | | |
| V13 | Vertical | Mono-evolved | | |
| V14 | Vertical | Mono-evolved | | |
| V15 | Vertical | Mono-evolved | | |

| | | | | |
|---|---|---|---|---|
| V16 | Vertical | Mono-evolved | | |
| H1 | Horizontal | Mono-evolved | 0.744 | ns |
| H2 | Horizontal | Mono-evolved | | |
| H3 | Horizontal | Mono-evolved | | |
| H4 | Horizontal | Mono-evolved | | |
| H5 | Horizontal | Mono-evolved | | |
| H6 | Horizontal | Mono-evolved | | |
| H7 | Horizontal | Mono-evolved | | |
| H8 | Horizontal | Mono-evolved | | |
| H9 | Horizontal | Mono-evolved | | |
| H10 | Horizontal | Mono-evolved | | |
| H11 | Horizontal | Mono-evolved | | |
| H12 | Horizontal | Mono-evolved | | |
| H13 | Horizontal | Mono-evolved | | |
| H14 | Horizontal | Mono-evolved | | |
| H15 | Horizontal | Mono-evolved | | |
| H16 | Horizontal | Mono-evolved | | |

**Supplementary Table T2:** Shows the statistically significant differences in the survival evolved *E. coli* from both the treatments when cocultured with ancestor *M. xanthus*. Letters indicate significant differences.

**Supplementary Table T3 (Attached excel file named *E.coli*_genomics.xls):** Shows the list of unique mutations only present in the evolved *E. coli* genomes from both coevolved and mono-evolved treatments. The prey clones good at resisting predators are marked with "***", the intermediate ones with "**" and the ones bad at resisting predation are marked with "*" (Attached excel file named *E.coli*_genomics.xls).

**Supplementary Table T4 (Attached excel file named *M. xanthus*_genomics.xls):** Shows the list of unique mutations only present in the evolved *M. xanthus* genomes from both coevolves and mono-evolved treatments relative to their ancestors. The clones that are bad at killing preys have been marked (Attached excel file named *M.xanthus*_genomics.xls).
